# The source of microbial transmission influences niche colonization and microbiome development

**DOI:** 10.1101/2023.01.31.526426

**Authors:** Isabel S. Tanger, Julia Stefanschitz, Yannick Schwert, Olivia Roth

## Abstract

Early life microbial colonizers shape and support the immature vertebrate immune system. Microbial colonization relies on the vertical route via parental provisioning and the horizontal route via environmental contribution. Vertical transmission is mostly a maternal trait making it hard to determine the source of microbial colonization in order to gain insight in the establishment of the microbial community during crucial development stages. The evolution of unique male pregnancy in pipefishes and seahorses enables the disentanglement of both horizontal and vertical transmission, but also facilitates the differentiation of maternal vs. paternal provisioning ranging from egg development, to male pregnancy and early juvenile development. Using 16s rRNA amplicon sequencing and source-tracker analyses, we revealed how the distinct origins of transmission (maternal, paternal & horizontal) shaped the juvenile internal and external microbiome establishment in the broad-nosed pipefish *Syngnathus typhle.* Paternal provisioning mainly shaped the juvenile external microbiome, whereas maternal microbes were the main source of the internal juvenile microbiome, later developing into the gut microbiome. This suggests that stability of niche microbiomes may vary depending on the route and time point of colonization, the strength of environmental influences (i.e., horizontal transmission), and potentially the homeostatic function of the niche microbiome.

## Introduction

The coevolution of the host with its microbiome permitted the intimate physical integration of microbes shaping development, nutrition and digestion (1–4), but also influencing behaviour (5,6) and immune functions (7–10). Microbes may even provide their hosts with evolutionary novelties facilitating the co-option of new lifestyles and the colonization of new habitats (11). Often these microbial communities are highly species, tissue, and development stage-specific, others may form complex long-lasting interactions with their host (12,13). Acquisition of specific microbes can be of horizontal source (environmental or close contact with conspecifìcs), vertical through gestation or incubation, or a mix of both transmission modes (11,14–16). In horizontal transmission interests of hosts and microbe may frequently oppose each other fostering antagonistic interactions (17,18,19). This is in stark contrast to vertical transmission, where the transmission of the microbe is directly linked to the reproductive success of its host favouring cooperation (15,17,20–22) and ensuring consistent transmission of the same lineage of microbe across generations (23). For the host and its symbiont to evolve as a unit, the coexistence of the host-microbe association with matching host-microbe genotype on evolutionary timescales is a prerequisite (17,23). Vertical transmission not only facilitates coexistence across host generations (24,25) but the coevolution of host and microbe can evoke closer host dependence, as demonstrated over the loss of host fitness upon removal of vertically vs. horizontally transmitted symbionts (23). Even without close host dependence, as a temporary partner that only has a small transient contribution, a vertically transmitted microbe was proposed to be able to play out its evolutionary role (11).

To date, only a few studies have assessed the microbial communities of parents, gametes, and offspring in marine organisms in depth (26–36). Vertical transmission pathways are numerous and range from the deposition of microbes into the oocytes (26,37) or embryos (38,39), over smearing of microbes onto the egg during oocyte development and oviposition (40,41), to a variety of intimate parental-offspring interactions, e.g., pregnancy, birth and physical contact (10,42–47). The multitude of vertical transmission routes and the fact that most transmission routes are intermingled within the maternal line (but see: (29)) make the study of vertical transmission a demanding task hindering the disentanglement of synergistic, additive or antagonistic transmission dynamics and their impact on the host physiology.

Upon colonization, members of the microbiome build the first line of defence as microbial biofilm, preventing detrimental microorganisms from attachment, replication and colonization. Becoming a member via horizontal transmission, can thus be a challenging task given strong within host microbe-microbe competition (14,48–50). Early niche colonizers prime and boost the immature vertebrate immunity (51) becoming adjuvants to the immune system supporting host humoral and cellular immune defence and competing against potentially virulent microbes (52,53). The immune system can learn to differentiate between friend and foe and develop in order to maintain a symbiotic relationship (54,55).

To gain insight into the development of the microbial community, we need to unravel the source of the microbial symbionts and assess niche colonization and expansion during different developmental stages. Spotting early colonizers inherited from parents through vertical transmission routes is particularly crucial. The unique male pregnancy evolution in pipefishes, seahorses and seadragons (the syngnathids) offers a differentiation of the intermingled routes of vertical transmission. While syngnathid females may deposit microbes into the oocytes, males may contribute microbes to the next generation throughout the brood pouch (27). To unravel how the maternal vs. the paternal routes of vertical transmission, as well as the horizontal route, drive embryonal and juvenile microbiome development, we conducted two experiments with the broad-nosed pipefish *Syngnathus typhle* spanning from egg development throughout male pregnancy and ending with the first days post juvenile release. These two experiments were conducted to gain insight into the nature of the maternally and paternally inherited microbes, the horizontal contribution and the microbiome establishment during the first few days of juvenile development. The first experiment aimed to identify differences in microbial community composition in both α-diversity and β-diversity, as well as defining key microbes and tracking microbial source before fertilization, during male pregnancy and during development in the paternal brood pouch. To shed light on the route of vertical transmission and to determine how the route of microbial transmission (maternal, paternal and horizontal) shapes microbiome establishment in distinct offspring niches, we sampled unfertilized eggs (surface sterilized or unmanipulated), male and female hindguts, male brood pouches, testes and embryos (surface sterilized and unmanipulated) at three time points during the pregnancy for microbial 16rRNA genotyping including environmental samples. Through the direct comparison of internal (surface sterilized) and external microbial community in both unfertilized eggs and developing embryos, we could elucidate the origin of organ specific microbial communities during embryogenesis and pregnancy and their contribution to the establishment of the internal vs. external microbiome.

In the second experiment, we assessed the microbial community of juvenile broad-nosed pipefish during the first 12 days post-release (dpr) from the brood pouch by calculating the same above-mentioned diversity indexes and indicator species analyses. Inferring about gut vs. whole body microbiome, this experiment illuminated the development of the microbiome from birth throughout the first environmental microbial contact through swimming and feeding, highlighting how the supposedly maternally and paternally vertically transmitted microbes play out when horizontal transmission routes take over in establishing the pipefish microbiome.

This study permits to pinpoint specific microbial genera in the parental organs, throughout different phases of pregnancy, and assess their contribution to offspring microbial colonization. The insights will advance our knowledge about how distinct routes of vertical transmission interact in microbial colonization and development. This will permit the initiation of experimental microbial manipulation in *S. typhle* with the aim to investigate the role of microbes in offspring development and immunity, and ultimately enlighten how microbes influence the unique lifestyle evolution in male pregnant syngnathids.

## Methods

### Sample collection

Adult *Syngnathus typhle* were caught in Orth on Fehmarn (54°26’N 11°02’E) and brought to our aquaria facilities at GEOMAR Kiel, for breeding. Fish were kept in a flow-through aquaria system at 18°C with 18h day/6h night light regime and fed with live *Artemia salina* and frozen *Mysidae* spp. twice a day. In each aquarium three males and three females were kept together to allow mating. After the onset of breeding, fish were randomly sampled, regarding their sex and gravity stages, on five days between the end of May 2019 and end of June 2019. We sampled 89 mature *S. typhle* (18 females, 16 early pregnant males, 19 mid pregnant males, 18 late pregnant males and 19 non-pregnant males). Pregnancy stages (early, mid and late pregnancy) were defined according to (56). In order to detect microbial transfer from parental gonads and pouch tissue to the juveniles, we sampled testes and endometrial inner pouch lining tissue as well as fertilized larvae from the three pregnancy stages in male fish. In female fish, we sampled unfertilized eggs to assess potentially deposited microbes into the eggs or on their surface. The hindgut was sampled irrespective of sex. To investigate maternal microbial transfer through the cytoplasm, we surface-sterilized half of the unfertilized eggs from each female in a bath of 0.5% Polyvinylpyrrolidone-iodine (PVP-I, Solution in sterile-filtered Phosphate buffered Saline (PBS)) for 5 min with subsequent washing three times with 500μl sterile-filtered PBS (adapted from (57)). To sample the cytoplasm of surface-sterilized eggs (sterilized eggs) without contamination through the chorion, the egg was squished in the collection tube. The same sterilization treatment was applied to larvae (sterilized juveniles) of different pregnancy stages to discriminate between external and the internal microbiome. Non-sterilized eggs (untreated eggs) and larvae (untreated juveniles) were directly placed in the collection tubes. We pooled three juvenile and eggs sample from each of the pregnancy stages and sterilization treatment. All sampled organs were collected into collection microtubes from the DNeasy96 Blood and Tissue Kit from Qiagen (Hilden, Germany) and stored immediately at −80 °C.

### Juvenile microbiota development

Pregnant male *S. typhle* were caught in Orth on Fehmarn (54°26’N 11°02’E) in late spring 2019, transferred to the aquaria system, kept individually in a flow-through system with 18h day/6h night cycle and fed twice a day with live *Mysidae* spp. After parturition, free-swimming juveniles of each male were kept in a distinct aquarium and fed *ad libitum* twice per day with live *Artemia salina.* First sampling took place after release from the brood pouch, sampling was then continued in 2-3 day intervals. During each sampling three juveniles per family tank were collected individually in a collection tube for the analysis of whole-body microbiome (whole juveniles) development. To test for development of internal *vs.* external microbiome another three juveniles from the same family tank were euthanized by immersion in MS222 and the gut was removed in a sterile manner (juvenile gut) and collected individually. At each sampling day, controls of water and food samples were taken.

### RNA extraction, library preparation and amplicon sequencing

Both datasets have been treated with the same DNA extraction and 16S rRNA sequencing protocols. DNA extraction was done with DNeasy Blood & Tissue Kit (QIAGEN, Germany) following the manufacturers protocol including a pre-treatment for Gram-positive bacteria with ameliorations from (58). See Supplementary Material 1 for further details. Library preparation was done by the institute for experimental medicine (UKSH, Campus Kiel) with 20μl of sample DNA from each sample. Amplicons of the V3-V4 hypervariable region (341f/806r) were sequenced using the Illumina MiSeq platform (Illumina, USA) with 2x 300-bp paired-end read settings at the IKMB, Kiel University.

### Data analysis

Demultiplexed sequences were processed using DADA2 implemented in the Qiime2 platform (version 2021.8(59)) for primer cutting, timing, quality control, merging, chimera removal and denoising. Taxonomy was assigned using the Silva 132 classifier for Qiime 2 (Version 2019.10) for the V3/V4 hypervariable region. Mitochondrial and chloroplast sequences were removed before further analyses. Further sorting and statistical analysis of exported Amplicon sequence variants (ASV) were conducted in R (Version 4.1.0, (60)) using the phyloseq package (Version 1.36.0,(61)). The two experiments were analysed separately using the same parameters. After removal of ASV’s with non-definite taxonomic classification (NA) in phylum, family and genera, we applied a prevalence filter of 2% and agglomerated the sequences on genus level.

FaithPD represents α-diversity (“microbiome”, version 1.14.0, (Lahti et al.,2012-2019)). Hypothesis testing using an aov (“stats”, version 4.1.0, (60)) with either blocked ANOVA for the adult treatment (x ~ gravity + organ, data =adult) or a repeated measures ANOVA for the juvenile data set (x~ treatmentAW*timepointAW, strata = familyAW, data = juvenile). In case of significant effects in the ANOVA a TukeyHSD test using false-discovery rate was applied as a post-hoc test.

β-diversity was tested on Bray Curtis dissimilarity matrix (BCdM) and UniFrac distance matrix (UFdM). The first calculates presence and absence and infers information about the numerical composition within / between microbial communities, the latter includes phylogenetic distance between the ASV and thus provides insight into the phylogenetic spread of a microbial community. β-diversity was tested in a two factorial blocked PERMANOVA (“vegan” version 2.5.7, (63)) (vertical transmission; adonis(x_brayCurtis~organ+gravity, data= adult, permutations 10000)) or a two factorial repeated measures PERMANOVA (juvenile development; adonis (x_brayCurtis~ treatmentAW* timepointA, data= juvenile, strata=familyAW, permutations 10000)) on BCdM and UFdM. Pairwise.adonis2 (version 0.4, (64)) was used as a post-hoc test to detect pairwise differences. False discovery rate was used for p-value adjustment to multiple testing. Data were visualized using a principal coordinate analysis of the 50 most abundant ASVs including 95% confidence ellipses.

An indicator species analysis (multipatt (“indicspecies”, version 1.7.9, (65)) was run on both datasets independently to identify more abundant genera in either single levels of each factor or level combinations. To estimate the proportion of ASV originating from parental organs (vertical transmission) and from environmental factors (juvenile development), we applied Bayesian community-level microbial source tracking (sourcetracker2 1.0.1,(66)), defining parental organs and control samples as source and sterile and non-sterile juveniles as sink microbiomes.

## Results

### Vertical transfer of microbiota

During denoising and filtering 12 samples (#23 mid testes, #4 & #21 none testes, #14 early sterilized juveniles, #19 & # 48 early pregnant testes, 2 PVP-I controls, #50 late pregnant sterilized juveniles, #54 sterilized unfertilized eggs, #70 mid sterilized juvenile, #74 non pregnant hindgut) were removed from the downstream analysis. After removing sequences with no taxonomic information (NA), 2% prevalence filtering and taxonomic agglomeration 278 unique genera have been identified. The most prevalent phyla were Proteobacteria (56.83%), Bacteroidetes (19.78%), Firmicutes (8.2%) and Actinobacteria (4.68 %).

Differences in phylogenetic diversity measured by the FaithPD existed between the organs irrespective of gravity (blocked ANOVA: Faith PD: organ: F_9_,355 = 6.741, p<0.001 ***, gravity: F_4,355_ = 0.611, p>0.05) (Figure 1). Sterilized eggs and sterilized juveniles had a lower phylogenetic diversity than untreated juveniles and the placenta. No differences in phylogenetic diversity were found between the untreated juveniles and the placenta or testes. Further, the placenta had a higher phylogenetic diversity than untreated eggs and the testes.

**Figure 1.**
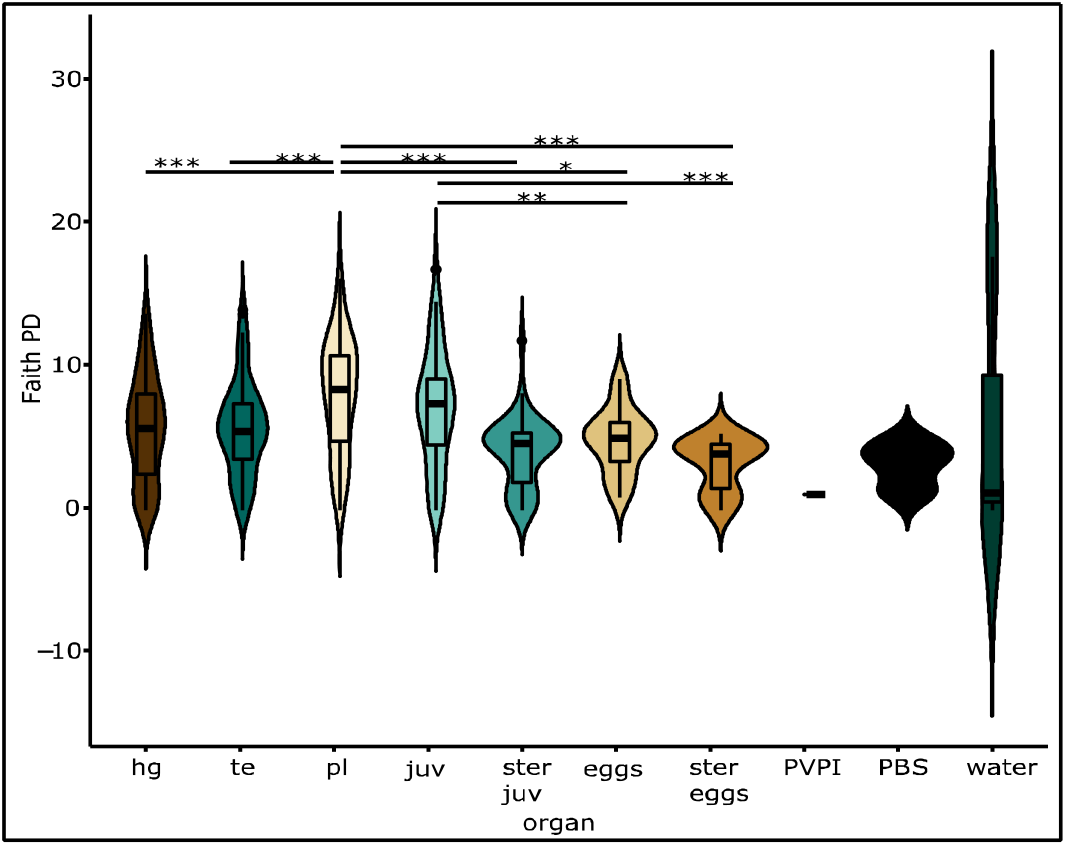
α-diversity index of organs important in parental transfer of microbiota in *S.typhle.* Violin plot of calculated Faith’s phylogenetic diversity. Boxes represent interquartile ranges between the first and the third quartile (25% and 75%) and horizontal lines define the median with a surrounding data probability density. Colours represent different organs, hg= hindgut, te= testes, pl= placenta, juv= untreated juveniles, ster juv= surface sterilized juveniles, eggs= untreated eggs, ster eggs= sterilized eggs, PVPI= Propidium Iodide, PBS= Phosphate buffered saline solution. Significant differences after post-hoc testing are shown using asterisks over connecting lines.

β-diversity, calculated from both the UFdM and the BCdM, differed for the organs (untreated juveniles, sterilized juveniles, untreated eggs, sterilized eggs, testes, placenta and hindgut) and gravity stages (female, non-pregnant, early, mid and late pregnant) (blocked PERMANOVA; Bray-Curtis dissimilarity matrix: organ: F_9,354_ = 6.13, p<0.001 ***, R^2^ = 0.13, gravity: F_4,354_ = 2.31 p<0.001 ***, R^2^ = 0.02; unweighted Unifrac: organ F_9,354_ = 4.65, p<0.001 ***, R^2^ = 0.1, gravity: F4,354 = 1.57 p<0.05 *, R^2^ = 0.01) (Figure 2). In the factor organ, pairwise comparisons of the UFdM revealed distinct microbiota composition between untreated juveniles and sterilized juveniles (p-value<0.01), untreated eggs (p-value<0.01), sterilized eggs (p-value<0.01) and water (p-value <0.05). However, a similar phylogenetic microbial setup was suggested in untreated juveniles and the placenta (p-value>0.05). Apart from this, all organs differed in their microbial composition. Pairwise comparisons of the BCdM, showed in addition to the aforementioned results a difference between untreated juveniles and the placenta (p-value placenta: untreated juveniles <0.01).

**Figure 2.**
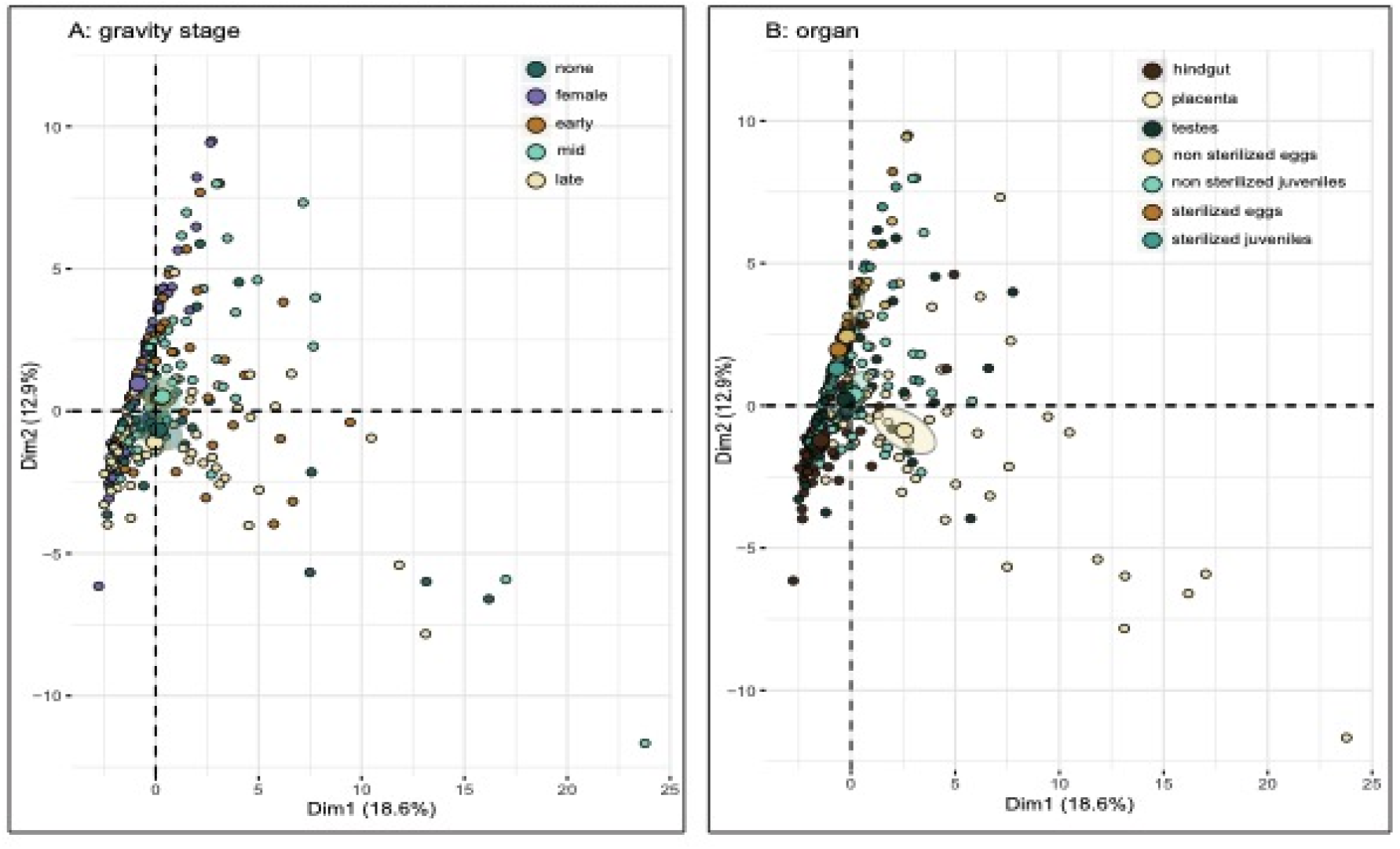
Principal component analysis (PCA) of the 50 most abundant microbial genera during *S. typhle* pregnancy. The first two principal components, Diml explaining 18.6% of the total variance and Dim2 explaining 12.9% of the total variance are shown together with 95% confidence ellipses (shaded area) around a centre of gravity (big points). Samples are indicated as small points. (A) Colours represent gravity stages associated top 50 microbial genera. (B) Colours represent organ association of top 50 microbial genera.

Effects of the gravity stage on the phylogenetically weighted β-diversity (UFdM) were found between females and late, mid and non-pregnant males (p-value female:late<0.05, f:mid<0.05, f:early<0.01). Additionally, late pregnant males differed from early pregnant males (p-value<0.01). The non-phylogenetically weighted BCdM also revealed differences between females and non-pregnant males (p-value 0.01) and late pregnant males (p-value<0.01) but also differences between late pregnant males and both early (p-value<0.01) and mid-pregnant males (p-value<0.01). For all statistical information see Supplementary Table 1.

Differences in the microbiome composition between organs or gravity stages were established with an indicator species analysis on all 278 genera. Details are presented in Supplementary Table 2 and Supplementary Table 3.

We could identify distinct groups of genera belonging to different organs. Specific groups were identified for the hindgut (4 genera)), untreated eggs (3 genera) and untreated juveniles (2 genera). Further, a large proportion of genera (18) was associated with the placenta. Untreated juveniles and the placenta shared 30 genera belonging mainly to the Proteobacteria and Bacteroidetes. Four genera associated with the male reproductive organs, testes and placenta all belonged to the Alphaproteobacteria. Unfertilized eggs, irrespective of their sterilization treatments were characterized by *Alishewanella. We* further identified a group of four genera *(Thalassotalea, OM27 clade, Octadecabacter, Olleya)* prevalent in untreated eggs, untreated juveniles and the placenta. 12 genera were associated with the male reproductive organs and untreated juveniles. Amongst which there were *Sulfitobacter* and *Lewinella. Granulicatella* was the only indicator shared by sterilized and untreated juveniles. Untreated eggs and untreated juveniles shared one genus of the *OM182* clade Gammaproteobacteria. Both genera, *Bacteroides* and *Pseudomonas* were prevalent in all organs and gravity stages assessed.

Indicator species analysis of the gravity stages has shown 12 genera associated with late pregnancy, belonging to the phyla Bacteroidetes and Proteobacteria, and three genera associated with non-pregnant males. Five genera in this group *(Aureispira, Ahrensia, Pseudahrensia,* uncultured *Flammeovirgaceae* and *Parvularcula)* were specific (A=1) to the male sex, these all belonged to either Alphaproteobacteria or Bacteroidia. In contrast, no genera specific to female organs were identified (sterilized and untreated eggs).

We conducted a source tracker analysis computing the proportion of juvenile microbiota (sterilized juveniles and untreated juveniles) originating from source microbiomes (all parental organs (sterilized eggs, untreated eggs, hindgut, placenta, testes) and the controls (PVP-I, PBS and water) (Figure 3). Proportions seen in Supplementary Table 4. Sterilized juvenile microbiome originated mostly from sterilized eggs, with some samples including also the hindgut, placental and untreated eggs. During male pregnancy, the overall picture of the distribution of source microbiome in the sterilized juveniles was rather consistent. In the untreated juveniles, the supposed source of the microbiome changed substantially over the course of pregnancy. In early pregnancy, results suggest that the microbiome was predominantly sourced from the sterilized eggs, however, this was in tandem with an increasing microbial contribution from the testes. Throughout pregnancy, the microbiome of the untreated juveniles changed from known sources of either sterilized or untreated eggs, placenta, testes and hindgut to a more unknown source of microbiome. In late pregnancy untreated juveniles, microbiome contributions from an unknown source was highest, followed by placenta, testes and untreated gonads. The proportion of microbes with a suggested source in seawater and the controls for PVP-I and PBS remained low in all samples.

**Figure 3:**
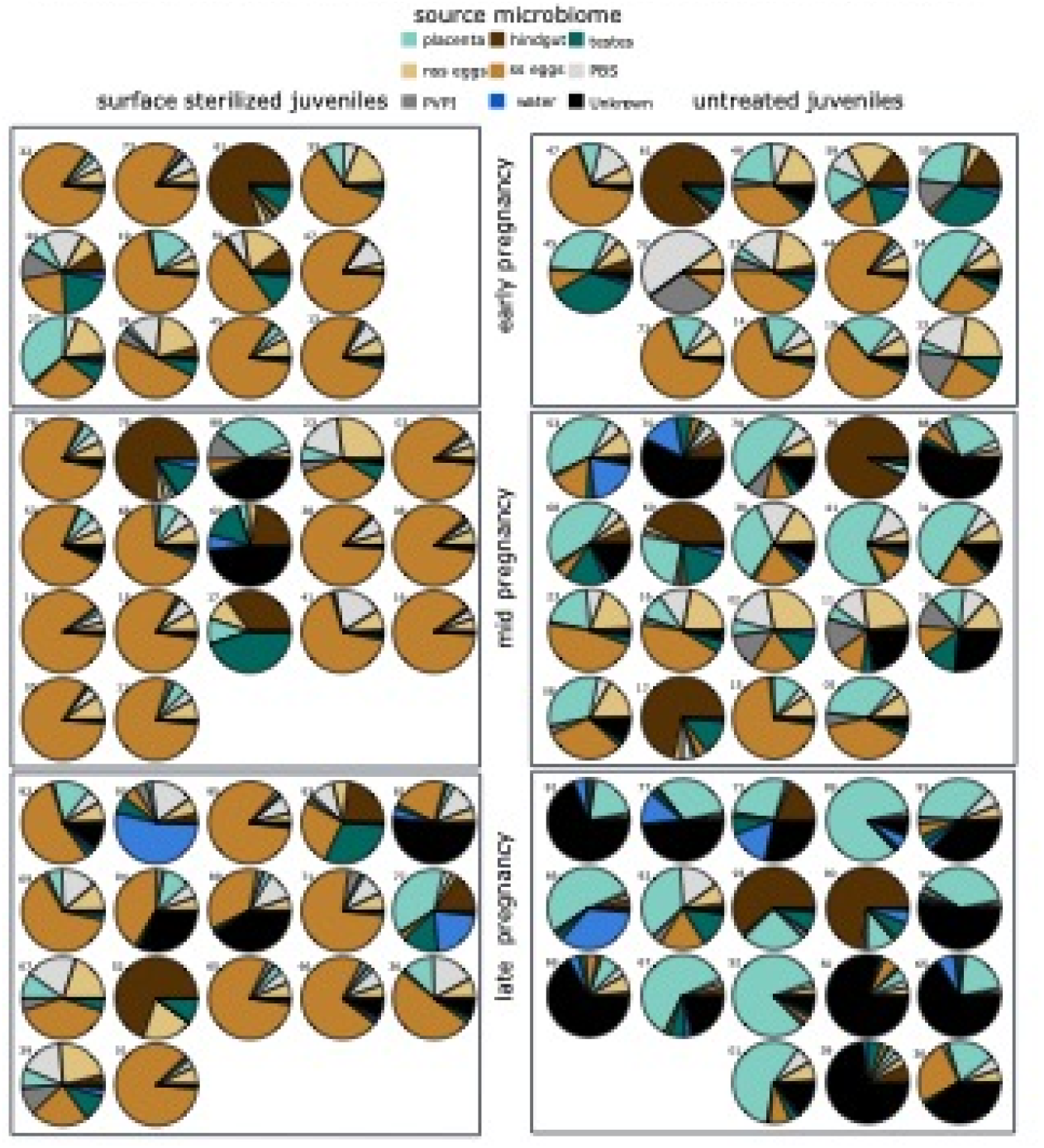
SourceTracker analysis of juvenile microbial community during pregnancy in *S. typhle.* Proportion of each source microbial community after 100 Gibbs samplings represented as partial circle in each sink sample. Sink samples were defined as untreated and surface sterilized juveniles during the different pregnancy stages (early, mid and late pregnancy). nss eggs= non-surface sterilized, untreated eggs/gonads and ss eggs= surface sterilized eggs/gonads

### Juvenile Development

To determine the course of development of the microbial community in juveniles up to 12 dpr, we sequenced 16S rRNA from whole body and gut of juveniles in interval of 2-3 days.

After removing sequences with no taxonomic information (NA), 2% prevalence filtering and taxonomic agglomeration at genera level we identified 495 unique genera. The most prevalent phyla over all samples were Proteobacteria (48.29%), Bacteroidetes (16.97%), Actinobacteria (6.27%) and Firmicutes (6.06 %).

### α-diversity

We used FaithPD as α-diversity measure between whole body and gut microbiota over sampling time. The repeated measures ANOVA (strata = family) showed an effect of parental origin on the treatment (whole body or gut) (repeated measures ANOVA FaithPD: treatment: F_1,351_ = 1314.708, p<0.001 ***, timepoint: F_6,351_ = 1.973, p>0.05 interaction terms: F_6,351_ = 3.341, p<0.01 **). FaithPD showed a higher phylogenetic diversity in whole-body juvenile microbiome compared to juvenile gut (p-value < 0.001), as well as differences in microbiomes of water and juvenile samples (p-value water-whole body < 0.01, water-juvenile gut<0.001) and differences between artemia and whole-body juveniles (p-value <0.001) (Figure 4). No significant differences in phylogenetic diversity of the sampled juvenile gut microbiome and artemia were detected (p-value >0.05). The phylogenetic microbiome diversity was not affected by sampling timepoint.

**Figure 4:**
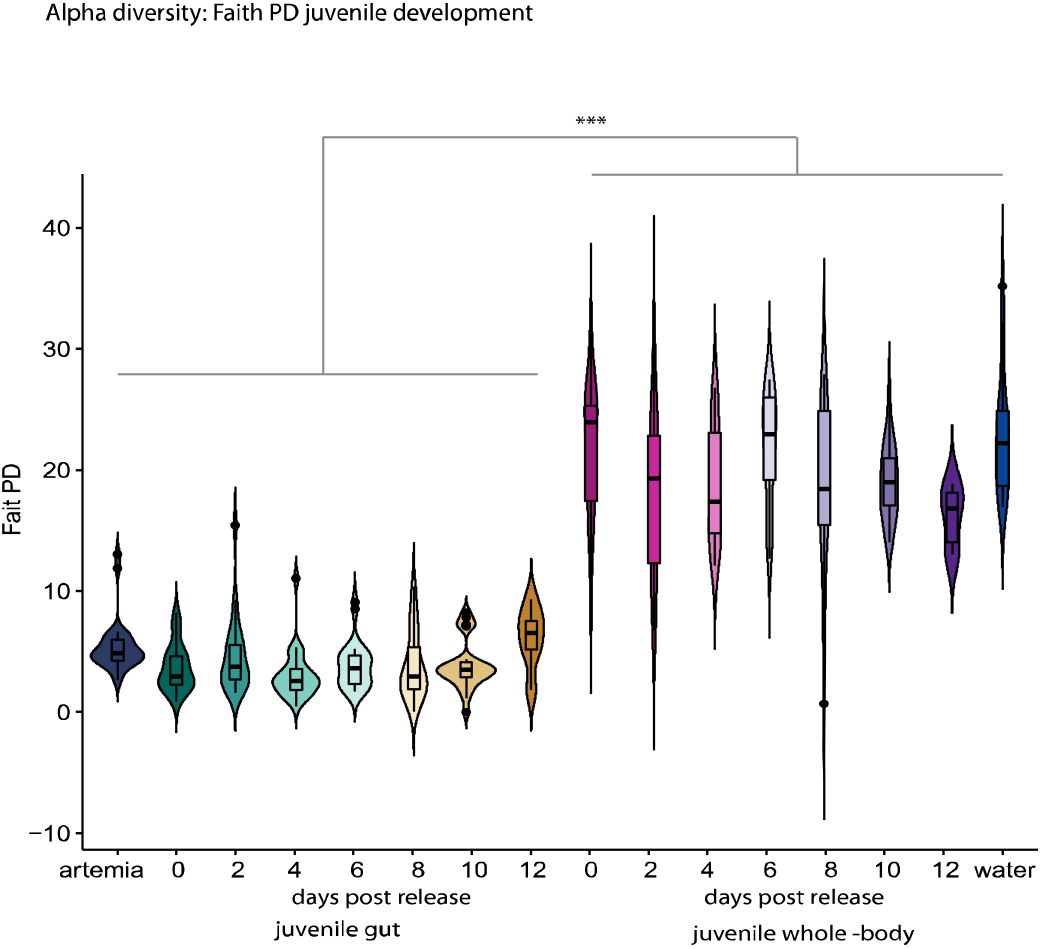
α-diversity index of juvenile whole body and gut microbial community development in *S.typhle.* Violin plot of calculated Faith’s phylogenetic diversity. Boxes represent interquartile ranges between the first and the third quartile (25% and 75%) and horizontal lines define the median with a surrounding data probability density. Colours represent different sampling timepoints and treatment. Significant differences after post-hoc testing are shown using asterisks over connecting lines.

### β-diversity

We computed a two-factorial repeated measures PERMANOVA on BCdM UFdM using timepoint of sampling and treatment as factors with family as strata. We identified differences in β-diversity between microbial communities in juvenile gut and whole-body juvenile over time in accordance with the influence of family on β-diversity of (repeated measures PERMANOVA: BCdM: treatment: F3,359 = 34.08, R^2^=0.2007, p<0.001 ***, timepoint: F_6,351_ = 5.146, R^2^=0.061, p<0.001 ***, interaction terms: F_6,351_ = 2,875, R^2^=0.034, p<0.001 ***; UFdM:: treatment: F_3,359_ = 73.198, R^2^=0.353, p<0.001 ***, timepoint: F_6,351_ = 4.276, R^2^=0.0413, p<0.001 ***, interaction terms: F_6,351_ = 2,785, R^2^=0.027, p<0.001 ***). According to post-hoc testing in BCdM, differences between whole-body juveniles and juvenile gut irrespective of the timepoint sampled were identified (p-value: whole-body (0dpr-12dpr):gut (0dpr-12dpr)<0.001). The microbial β-diversity of whole-body juveniles differed among all timepoints (p-value < 0.01, see supplementary Table 5) except from 2dpr to 4dpr (p-value>0.05), 4dpr-6dpr (p-value >0.05), 6dpr to 8dpr (p-value >0.05) and 8dpr-10dpr (p-value >0.05). The β-diversity of the juvenile gut microbiome differed between day of release and all other sampling timepoints (gut 0dpr: gut (2-12dpr) <0.01), further differences were found between 2dpr and 4dpr (p-value< 0,05), 2dpr and 6dpr (p-value <0.05) and 2dpr and 10dpr (p-value<0.01). Samples from 4dpr differ in their β-diversity from samples from 10dpr (p-value<0.01) and samples from 4dpr and 6dpr differed from 12dpr (4dpr: 12dpr: p-value<0.05, 6dpr:12dpr p-value<0.05). Additionally, β-diversity of juvenile gut microbiome between 8dpr to 12dpr differed (p-value<0.05). All samples, irrespective of timepoint and treatment had a distinct β-diversity than both water (p-value <0.01) and artemia (p-value <0.01) (Figure 5). Similar results were found in the post-hoc test of the phylogenetically weighted UFdM analysis. In contrast to the BCdM results, there were no significant differences in the whole-body juveniles between 2dpr and 4dpr (p-value<0.05) and between 4dpr and 6dpr (p-value<0.01). However, the juvenile gut microbiome seemed to be more constant over time. As such, 2dpr did not differ from any other timepoint except 10dpr (p-value < 0.05) and the microbial β-diversity of the 8dpr juvenile gut was similar to the microbial β-diversity of the 12dpr juvenile gut.

**Figure 5:**
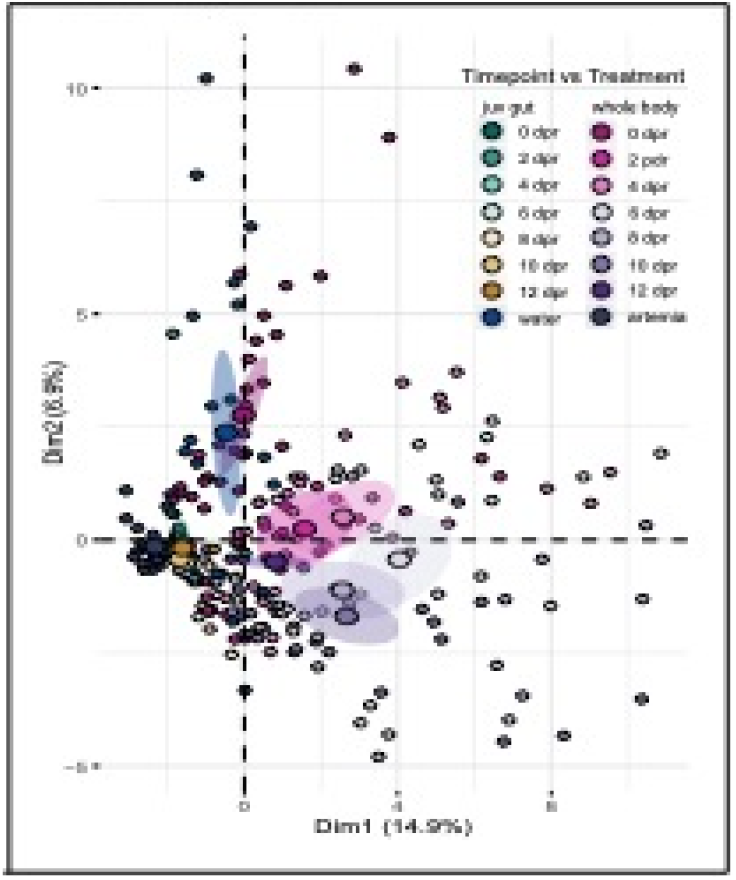
Overall principal component analysis (PCAļ of the 50 most abundant microbial genera during *S. typhle* juvenile development. The first two principal components, Dim1 explaining 14.9% of the total variance and Dim2 explaining 6.9% of the total variance are shown together with 95% confidence ellipses (shaded area) around a centre of gravity (big points). Samples are indicated as small points. colours represent treatment (juvenile gut or juvenile whole body microbial community) and timepoint of sampling as days post-release (dpr) associated top 50 microbial genera.

### Indicator species analysis

We could identify ten groups of genera indicative of water, artemia, whole body juveniles or juvenile gut or a combination of the four. The largest group was indicative of water samples which included 175 genera, further, we found indicator species of artemia (13 genera), whole body juveniles (24 genera) and juvenile gut (3 genera). 7 genera were specific for the whole-body microbiome of developing juveniles irrespective of timepoint and 34 genera specifically charactered water and whole-body juveniles. Within the indicator species group of the juvenile gut there were three species, *Yersinia, Pelomonas* and *Aeromonas* with a high specificity estimate (A> 0.95). Indicator species analysis of the timepoints irrespective of treatment was more diverse. Most genera were indicative of several timepoints and built an indicator group with either water or artemia. One genus, *Sandaracinus,* was specific to the intermediate time points 2dpr −12dpr. One genus (order Granulosicoccales) was specific for all timepoints except the release day. The genera specific to the water sample were the same as in the indicator analysis of the factor treatment. Results can be found in Supplementary Table 6 and Supplementary Table 7.

## Discussion

To unravel how the maternal *vs.* paternal routes of vertical transmission and the environmental contribution via the horizontal route drive embryonal and juvenile microbiome development in the broad-nosed pipefish *Syngnathus typhle, we* have assessed 16s rRNA microbial diversity in a range of maternal and paternal organs, and in different stages of the developing offspring, as well as in the environment.

In the first part, we have evaluated changes in microbial composition throughout reproduction and male pregnancy comparing the developing larval microbiome to the unfertilized maternal egg, the paternal testes and the paternal placenta-like tissue of *S. typhle* (1). To determine maternal microbial contribution, we compared the egg surface microbiota, the egg-internal microbiota and both surface sterilized and untreated larvae at three timepoints during pregnancy. To detect paternal origin, we compared placenta-like structure and testes of non-pregnant males to untreated larvae at three timepoints throughout the pregnancy. In addition, surrounding waters were assessed to understand the environmental (horizontal) microbial contribution to the offspring. In the second part, we have sampled juveniles (full sibs) over the first days post-release to understand microbial establishment (2). Here, we differentiated between the juvenile gut microbiome and the whole-body juvenile microbiome. In doing so, we aimed to resolve the development of the early microbiome in order to distinguish the importance of environmental and parental microbiota. In the third part, we have compared the indicator species analyses from both datasets to identify key microbial genera both during pregnancy and early microbiota development in free swimming juveniles. With this, we aimed to determine microbial genera that are shared between the juveniles during pregnancy and identify microbial genera associated with early development that should be studied in future microbial manipulation experiments (3).

### 1. Parental transfer of microbes

To assess offspring microbiome development across the gravity stages, microbial compositions in females, non-pregnant, early, mid and late pregnant males are discussed first disregarding organ specificity. While microbial α-diversity was not affected by gravity stage, β-diversity has shown both phylogenetic (UFdM) and compositional (BCdM) differences in microbial diversity. Non-pregnant males and females differed in their microbial composition indicating a sex-specific microbiome mainly driven by the reproductive organs (female gonads and male testes and placenta-like system) as no sex-specific effect was found in the hindgut. Late pregnant microbial composition differed from the early and mid-pregnancy indicating a shift in microbial composition over time. This shift supports previous insights into the parental microbiome of *S. typhle* (27) and matches a similar pattern in the human vaginal microbiome during the course of mammalian pregnancy (67). The transfer of eggs into the brood pouch influenced the microbial composition in the brood pouch, as indicated by a missing phylogenetic difference in the microbial community of female eggs and early pregnant males, in contrast to the phylogenetic microbial difference between female eggs and mid, non and late pregnant males. This suggests the transfer of maternal microbes to “feminized” the male brood pouch at the onset of the pregnancy, while subsequently the male microbiome was restored in mid and late pregnancy.

To gain insight into organ-specific microbes, we compared the microbial community composition across organs (placenta-like tissue, testes, hindgut, sterilized and unsterilized eggs, sterilized and unsterilized juveniles) irrespective of the stage of pregnancy. The placenta-like tissue has a higher α-diversity than any other organ except untreated juveniles (Figure 1). This high microbial diversity in the placenta-like structure as the central brooding organ, might be adaptive by protecting the highly vulnerable developing juveniles from potentially virulent microbes (68). This is essential at the onset of pregnancy when eggs are transferred into the brood pouch, but also during the last third of the pregnancy when the brood pouch becomes more permeable and thus sensitive to environmental influence (27). As the place of paternal vertical transfer, a diverse microbiome in the placenta-like tissue might be favourable for the unborn juveniles, permitting a diverse initial microbial colonization. Evidence for such vertical transfer can be found in the similar Faith PD Index and phylogeny incorporating UFdM of both placenta-like system and untreated juveniles hinting towards a phylogenetically similar microbiome in both organs (Figure 1 & 2B). Interestingly, there was no such grouping in the non-phylogenetically BCdM indicating similar ASVs with distinct numerical compositions in the placenta-like tissue and untreated juveniles. This suggests that paternally influenced juvenile microbial composition still needs to develop in terms of numerical proportions of microbial genera possibly influenced by the presence and abundance of maternally transferred ASV’s.

Maternal microbial transfer could be identified comparing untreated juveniles with both surface sterilized juveniles and sterilised/ unsterilised eggs. Considering, that mouth opening only occurs at the end of pregnancy (69), previous paternal vertical transfer to the internal microbiome of the juvenile is unlikely. Microbial communities identified in the surface sterilised juvenile might thus likely have maternal origin as a main source. The possibility of maternal transovarial transfer of the microbiome was supported by a similar microbiome in both untreated and sterilized eggs and sterilized juveniles as suggested in both α and β-diversity (Figure 1 & 2B). In sterilized eggs, the source tracker analysis identified a microbial contribution from natural sources that consisted through the pregnancy stages (sterilised and untreated eggs) (Figure 3). This permits to speculate that transovarial maternal microbes contribute to the initial gut microbiome before first feeding. Opposed to the rather constant microbiota composition of the surface sterilized juveniles, the microbiota of the untreated juveniles, i.e., the whole-body microbiome, underwent severe changes throughout pregnancy regarding the suggestive source of the microbiome. In early pregnancy larval stages, the contribution of sterile egg microbiota resembled those of the sterilized juveniles as a key microbiome source suggesting a delay in the paternal vertical transfer through the placenta at the onset of pregnancy. During the course of pregnancy, a shift in the source microbiome of the untreaded juvenile is indicated by rising proportion of paternally originated microbiome (placenta-like tissue) and decreasing influence of maternal source (eggs). Following the hatching of the pipefish embryos in the brood pouch (56), the paternal microbial influence on the juveniles must have increased (Figure 3), supporting our previous results (27). In late pregnancy, to accommodate for juvenile growth, the brood pouch becomes more permeable increasing the likelihood of horizontal transmission of the juvenile microbiome through environmental seawater. While the microbiota of untreated juveniles can be traced back to paternal organs as source microbiota in early and mid-pregnancy, untreated juveniles in late pregnancy exhibit a higher proportion of unknown microbial source probably of environmental origin indicating the development towards a parentally independent microbial community (Figure 3).

Our data suggests an important role of transovarial microbial transfer in the establishment of the internal juvenile microbiome potentially developing further into the gut microbiome. Paternally transferred microbial communities rather contributed to the surface external juvenile microbiome, potentially having a priming and protective effect on the unborn juvenile, with a gradually decreasing influence towards parturition.

### 2. Juvenile microbial community development

During pregnancy, the brood pouch microbiome, in tight interplay with the paternal immune system, is protecting the highly vulnerable juveniles from exposure to virulent infections. At birth, the release of the juveniles into the surrounding waters suddenly changes the requirements on the function of the juvenile microbiome. These environmental microbes will from now on be the main source of colonization and are supposed to shape the offspring microbiome and immune system. The second part of our study provides insights into the gut vs. the whole-body microbiome of pipefish juveniles over the first 12 days post-release from the brood pouch.

Already in freshly released juveniles, the internal gut microbiota and whole-body microbiota can be clearly distinguished (as indicated by the α-diversity and the β-diversity) (Figure 6B). This suggests that the priming effect provided by the vertically transmitted maternal and paternal microbes was essential for early microbial niche colonization permitting the establishment of an initial gut microbiome in *S. typhle* juveniles.

**Figure 6:**
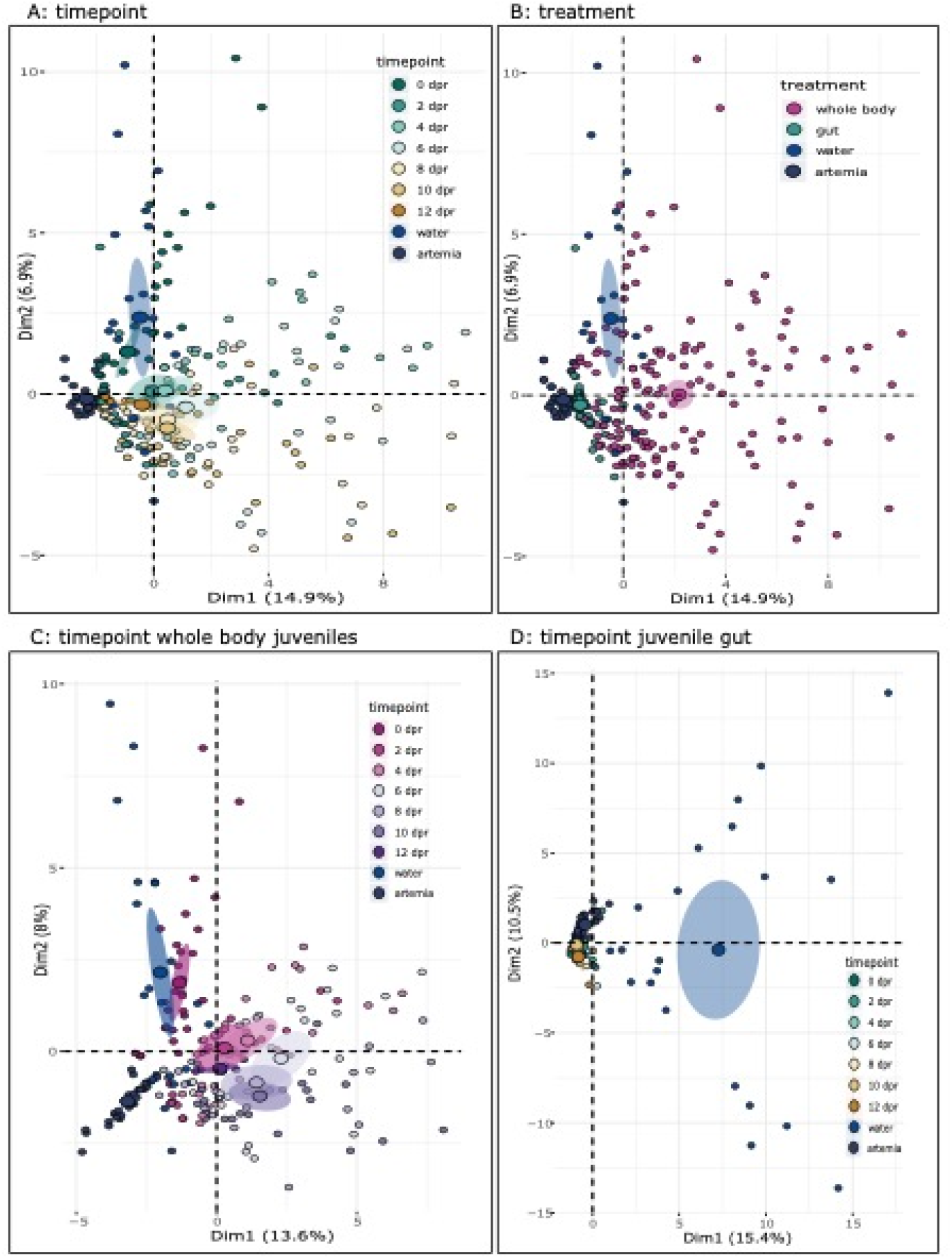
Specific principal component analysis (PCA) of the 50 most abundant microbial genera during *S. typhle* juvenile development. Included are the first two principal components of the top 50 most abundant ASV together with 95% confidence ellipses (shaded area) around a centre of gravity (big points). Samples are indicated as small points. (A) & (B) Dim1 explaining 14.9% of the total variance and Dim2 explaining 6.9% of the total variance. (A) PCA of the effect of temporal change on the top 50 most abundant microbial genera during *S. typhle* juvenile development irrespective of treatments. (B) PCA of the effect of sampled organ, juvenile whole body or juvenile gut, on the top 50 most abundant microbial genera during *S. typhle* juvenile development irrespective of sampling timepoints. (C) PCA of the temporal change on the top 50 most abundant microbial genera in juvenile whole-body microbiome. Dim1 explaining 13.6% of the total variance and Dim2 explaining 8% of the total variance. (D) PCA of the temporal change on the top 50 most abundant microbial genera in juvenile gut microbiome. Dim1 explaining 15.4 % of the total variance and Dim2 explaining 10% of the total variance.

The β-diversity of the gut microbiome of newly released juveniles was less variable and has shown only little changes over early juvenile development compared to the whole-body microbiome (Figure 5 and Figure 6B). Statistically, most of the differences in the interaction term could be assigned to the substantial changes of the whole-body microbial community changes during early free-living development (Figure 6C, Figure 6D), indicating a higher influence of horizontally transferred microbiota on the whole-body microbiome during the first few days after paternal release.

In the whole-body juvenile microbiome, we could detect two critical microbial shifts after parturition (Figure 6C). The first shift in β-diversity occurred between 0 and 2 dpr, supposedly imposed by the first contact to seawater and the start of feeding. At this stage, the microbiome of the whole-body juvenile was, in addition to the vertically transmitted genera, further enriched by horizontally transmitted microbes of food and seawater. The second shift occurred between 2 & 4 dpr and 6, 8 & 10 dpr. Cyanobacteria and Acidobacteriota were key bacteria groups defining the microbiota community of freshly released juveniles (Supplementary Table 7). In contrast, the juvenile microbiome 6-10 dpr was defined by the genus *Rubitalea* from the phylum Verrucomicrobiota, interestingly, a genus also abundant in the hindgut of adult *S. typhle* and in untreated juveniles during pregnancy. An acclimatization phase of the internal microbiome to environmental microbes (food and water) could explain why *Rubitalea* is present in the juvenile during pregnancy but not in the first dpr. The decrease in intraindividual variation of the whole-body microbiome at 12 dpr is explained by the establishment of a stable core microbiome as represented by a set of microbial taxa shared by most *S. typhle,* and supported by cases in the human microbiome (70). A core microbiome can be temporally induced and remain stable over a certain life stage such as e.g., pregnancy or juvenile development (71). Insights into temporal core microbiomes will facilitate further investigations about the function of certain microbes ultimately permitting the identification of the host-adapted core microbes influencing host fitness (72) that have a high probability to be vertically transmitted (73).

### 3. Early life key microbial communities

We aimed to identify key microbial genera in *S. typhle* development that are important in the establishment of a stable microbiome. These will be candidate genera for future manipulation experiments permitting the identification of their functions in male pregnancy, juvenile development and immune system maturation. To identify key microbial genera, we conducted four indicator species analyses (one for each factor in both data sets; adult organ (male and female reproductive organs, sterilized and untreated juveniles), adult gravity (females or male pregnancy stages), juvenile treatment (gut or overall microbiome) and juvenile timepoint (0 −12 dpr, water and artemia control) over all genera identified in both experiments (vertical transfer and juvenile development). We searched for genera present in two or more indicator species analyses. We have identified three bacterial genera *(Corynebacterium, Streptococcus, Yersinia)* present in the gut of free-swimming juveniles and either treated eggs or juveniles (Table 1). One of these, *Corynebacterium* was specific for treated eggs and the juvenile gut and is thus a strong candidate for transovarial transfer to the juvenile gut microbiome. The other two, *Streptococcus* and *Yersinia* were indicator genera of sterilized eggs and sterilized juveniles, and the juvenile gut microbiota, they were also present in the placenta-like tissue and the testes as well as in untreated eggs and juveniles.

**Table 1:**
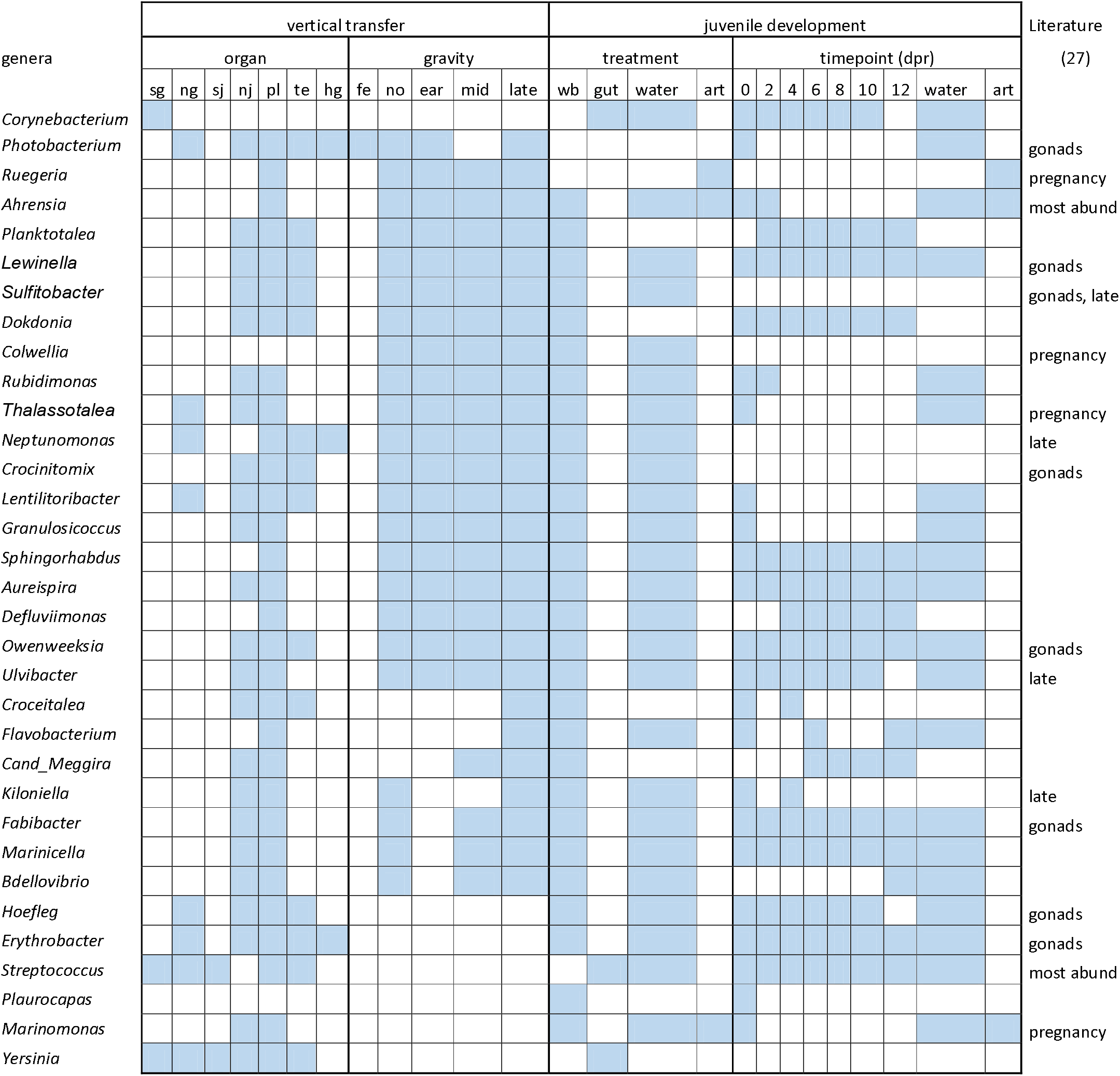
Key Microbial genera for vertical transfer and juvenile development: Microbial genera present in more than two indicator species analysis (multipart) of microbial community of either parental organs and parental gravity stages or juvenile treatment and timepoint in comparison important microbial genera identified in previous studies. Abbreviations used are hg = hindgut, te = testes, pl = placenta, nj = untreated juveniles, sj = surface sterilized juveniles, ng = untreated eggs / female gonads, sg = sterilized eggs / female gonads. Fe = female, no= none pregnant, ear=early pregnant, wb= whole-body, art = artemia

We suggest *Planktotalea, Dokdonia, Croceitalea* and *Candidatus Megaira* as key genera for paternal microbial transfer, as they were indicator species of the untreated juveniles and the placenta, as well as of the whole-body microbiome of the juveniles. Genera indicative of both male reproductive organs and untreated juveniles and whole-body juveniles could serve as candidates for a potential *S. typhle* core microbiome. Of special interest are nine of these 17 genera *(Lewinella, Sulfitobacter, Thalassotalea, Neptunomonas, Crocinitomix, Owenweeksia, Ulvibacter, Kiloniella* and *Fabibacter),* which have already been identified as potential candidates for vertical transmission (27). Most bacterial genera with an involvement in paternal transfer to untreated juveniles and the whole-body microbiome of free-swimming juveniles were present in all pregnancy stages and were identified in most timepoints in free-swimming juveniles. *Croceitalea* and *Kiloniella,* however, were only indicative of late pregnancy and during the first few sampling timepoints of free-swimming juveniles, rendering them candidates for primary horizontal colonizers of the juvenile microbiome. An additional group of bacterial genera *(Sphingorhabdus* and *Defluviimonas)* were indicative of the placental microbiome in either all pregnancy stages or only the late pregnancy stage *(Flavobacterium),* as well as the whole-body microbiome of free-swimming juveniles. Their presence in the placenta and their absence in untreated juveniles suggests a potential colonization during juvenile release from the brood pouch. In comparison with previous studies, two microbes *(Ruegeria* and *Ahrensia)* have been already proposed to be possible candidates for parental microbial transfer in *S. typhle* (27). Here, in contrast, they were not only indicative of the placenta in each pregnancy stage but were associated with both artemia or seawater and whole-body microbiomes (Table 1)

The close contact of pipefish juveniles with their father over the placenta-like system during male pregnancy provided an opportunity for the evolution of bi-parental trans-generational microbial transfer. In this study, we provide evidence for distinct routes in maternal *vs.* paternal transmission of microbes to pipefish juveniles that potentially provide synergistic effects. We highlight that the paternal influence shapes predominantly the external microbiome of the juvenile during male pregnancy. In contrast, we suggest transovarial transferred maternal microbes as a main source of the internal juvenile microbiome that eventually establishes the gut microbiome. The maternally transferred microbes established a stable community that persisted also in the gut of free-swimming juveniles suggesting early transovarial transmission to provide essential microbes with long-lasting influence in life of pipefish juveniles. Their early transmission as well as their colonization of internal niches may have provisioned the transovarial transmitted microbes with an advantage against later colonizers due to strong competitive abilities and their hiddenness from subsequent environmental influences. We thus suggest colonization of niches and the subsequent stability of niche microbiomes to depend on the route of transmission, the timepoint of colonization and the susceptibility to environmental influences (including the impact of horizontal transmission). Evaluating the combination of these factors separately for multiple intermingled but potentially synergistic routes of transmission will facilitate the assessment of host colonization and microbiome establishment, and ultimately foster our understanding of what defines differences in homeostatic functions across distinct niche microbiomes.

Future studies should restrict sampling to the here suggested crucial timepoints, and instead, follow the juvenile development post paternal release for a longer period. Further, the dissection of the juvenile gut should be established already during pregnancy, to enable an earlier differentiation between the gut and the whole-body microbiome providing a more detailed insight into gut microbiota establishment and the specific impact of maternal vertical microbiota transfer. Altogether, this will provide a higher resolution of the maternal *vs.* paternal role in microbial transfer in contrast to the horizontal transfer and on the development of the juvenile microbiome. To disentangle parental (maternal *vs.* paternal) from horizontal microbial colonization in more depth and follow niche colonization and microbe-microbe competition, we require data of the parental, larval and juvenile from several pipefish generations in order to find persistent members of the core microbiome. For a detailed investigation of the microbial colonization of pipefish eggs, larvae and juveniles, and an assessment of the functions of key microbial species strains, the time for experimental microbial manipulation of the maternal and paternal microbiome has arrived. Tracing key microbes from the parents to the offspring through genetic fluorescent marking will ensure their route of vertical *vs.* horizontal transfer. By removing and adding bacterial strains and investigating their physiological impact on the host-side, we can bridge the route of transfer to other physiological traits of the pipefish life, such as their development and their immune system.

## Acknowledgements

We thank Silke-Mareike Maarten, Diana Gill and Katja Cloppenborg-Schmidt for their support in the laboratory and for library preparation, Fabian Wendt and Johannes Hasse for fish tending and aquaria facility maintenance, Sören Franzenburg and his team for Illumina MiSeq Sequencing. We want to thank Arseny Dubin for general data curation and help during data analysis and Jamie Parker for proof-reading of this manuscript. We are grateful for the never-ending support by Thorsten Reusch who has hosted the pipefish group from 2009-2021 at GEOMAR, it was a blast. We thank the MarEvol group and members of the CRC 1182 Origin and Function of Metaorganism for fruitful discussions creating an excellent host-microbe environment at Kiel University.

## Author contribution

OR and IST have initiated and planned this project; IST and JS have conducted the experiments and the laboratory work; IST and YS have analysed the data. IST and OR have interpreted the data and written this manuscript.

## Supplementary Material

Supplementary Table 1: β-diversity vertical transfer

Supplementary Table 2: Indicator analysis vertical transfer; organs

Supplementary Table 3: Indicator analysis vertical transfer; gravity stages

Supplementary Table 4: SourceTracker proportions

Supplementary Table 5:: β-diversity juvenile development

Supplementary Table 6: Indicator analysis juvenile development; treatment

Supplementary Table 7: Indicator analysis juvenile development; timepoints

Supplementary Material 1: DNA Extraction Protocoll

