## Supplementary Material 1 for "The source of microbial transmission influences niche colonization and microbiome development"

Supplementary Matrial 1:

DNA Extraction

Both datasets have been treated with the same RNA extraction and 16S rRNA sequencing protocols. DNA extraction was done with the DNeasy Blood & Tissue Kit (QIAGEN, Germany) following the manufacturers protocol including a pre-treatment for Gram-positive bacteria with ameliorations from (1). The collected tissue was lysed in 180µl lysozyme solution (20mM Tris CL pH 8.0, 2mM sodium EDTA, 1.2% Triton X-100, lysozyme 20mg/ml) at 37°C for 2 hours on the shaker. For proteinase K digestion 25µl of proteinase K and 200µl of buffer AL (without EtOH) was mixed, added to the sample and incubated at 56°C. After 1h 200µl 99%-100% EtOH was added and the sample was mixed by vortexing. gDNA extraction was done following the manufacturers protocol with an overnight lysis step. After lysis and two washing steps DNA was eluted from the column with double elution step with 60µl buffer AE each

1.	Korsch M, Marten SM, Walther W, Vital M, Pieper D, Dötsch A. Impact of dental cement on the peri-implant biofilm-microbial comparison of two different cements in an in vivo observational study. Clin Implant Dent Relat Res. 2018 Aug 20;20.
